# Modeling sound attenuation in heterogeneous environments for improved bioacoustic sampling of wildlife populations

**DOI:** 10.1101/239079

**Authors:** J. Andrew Royle

## Abstract

Acoustic sampling methods are becoming increasingly important in biological monitoring. Sound attenuation is one of the most important dynamics affecting the utility of bioacoustic data as it directly affects the probability of detection of individuals from bioacoustic arrays and especially the localization of acoustic signals necessary in telemetry studies. Therefore, models of sound attenuation are necessary to make efficient use of bioacoustic data in ecological monitoring and assessment applications. Models of attenuation in widespread use are based on Euclidean distance between source and sensor, which is justified under spherical attenuation of sound waves in homogeneous environments. In some applications there are efforts to evaluate the detection range of sensors in response to local environmental characteristics at the sensor or at sentinel source locations with known environmental characteristics. However, attenuation is a function of the total environment between source and sensor, not just their locations. In this paper I develop a model of signal attenuation based on a non-Euclidean cost-weighted distance metric which contains resistance parameters that relate to environmental heterogeneity in the vicinity of an array. Importantly, these parameters can be estimated by maximum likelihood using experimental data from an array of fixed sources, thus allowing investigators who use bioacoustic methods to devise explicit models of sound attenuation *in situ*. In addition, drawing on analogy with classes of models known as spatial capture-recapture, I show that parameters of the non-Euclidean model of attenuation can be estimated when source locations are *unknown*. Thus, the models can be applied to real field studies which require localization of signals in heterogeneous environments.

## 1 Introduction

Acoustic sampling technology has emerged as an important method for the study of vocal species such as birds, anurans, marine mammals, many species of fish, and primates, or for species to which acoustic transponders can be implanted or affixed to. As a result, the deployment of automated acoustic recording devices has proliferated rapidly in both terrestrial (Blumstein et al. 2011; Digby et al. 2016; Brauer et al. 2016; Measey et al. 2017) and aquatic (Marques et al. 2009; Kessel et al. 2013; Marques et al. 2013; Cooke et al. 2016; Crossin et al. 2017) systems.

Bioacoustic technology is broadly relevant to the study of spatial ecology of animal populations. Two specific uses which are the focus of this paper are the application of bioacoustics to density estimation using variations of spatial capture-recapture (SCR) methods (Efford et al. 2009; Marques et al. 2013; Stevenson et al. 2015; Kidney et al. 2016) and its use in acoustic telemetry (Heupel et al. 2006) for the study of movement and resource selection. Acoustic telemetry has become widely used in aquatic environments to study fish, sea turtles, and marine mammals. Use of acoustic data for either SCR or telemetry requires *localization* of the observed signals obtained from acoustic data. This is essentially statistical triangulation which can be done when signals are obtained from an array of bioacoustic sensors so that potentially multiple detections of the same signal are possible (Janik et al. 2000; McGregor et al. 1997; Bower and Clark 2005; Blumstein et al. 2011). The precision of source localization improves with the number of sensors in the array and the density of the array. Localization has been recognized as being analogous to inference about the activity center in SCR methods, and therefore SCR has been adapted to accommodate data obtained by acoustic sampling methods (Dawson and Efford 2009; Efford et al. 2009; Borchers et al. 2015; Stevenson et al. 2015; Kidney et al. 2016).

Localization of acoustic sources requires explicit models for sound attenuation, i.e., the energy loss of sound propagation through a medium. In general, attenuation depends on the properties of the medium (Wiley and Richards 1982), and this often is characterized experimentally by engineers to satisfy design objectives of acoustic systems. However, to date, applications of bioacoustic methods in ecology have used simplistic models of spherical attenuation, in which amplitude decays according to a power law with rate proportional to the inverse of Euclidean distance^1^. In practice, sound attenuation is strongly affected by the structure of the environment between the source origin and the receiver (Singh et al. 2009; Kessel et al. 2013; Rek and Kwiatkowska 2016; Selby et al. 2016). When the environment is highly heterogeneous Euclidean distance models may be inadequate. For example in bird monitoring problems there may be substantial variability in vegetation density or height in the vicinity of a sensor or array of sensors. In acoustic telemetry studies, the attenuation of the signal can depend on depth, substrate, surface conditions and many other factors (Selby et al. 2016). This has led to considerable recent attention to the problem of “range testing” to determine effective detection range given environmental heterogeneity for acoustic telemetry applications (Marques et al. 2009; Kessel et al. 2013; Selby et al. 2016). For example, Selby et al. (2016) model attenuation as a function of source and sensor specific covariates (e.g., depth). However, attenuation of signals depends on the total environment between the signal and the source and therefore more general models of attenuation are needed.

Ideally, the end use of bioacoustic data in monitoring and assessment of biological populations should integrate explicit models of sound attenuation with parameters that are themselves estimated *in situ* along with biological parameters of interest such as density, position of sources, occupancy, or other ecological state variables. In this paper I suggest flexible classes of models for modeling attenuation in heterogeneous environments using cost-weighted distance in which effective distance is defined by a cost function that involves spatially explicit structure describing a heterogeneous landscape. This non-Euclidean distance model is widely used in least-cost path model analysis (Adriaensen et al. 2003) of landscape connectivity. Inference under this model has been formalized in the context of spatial capture-recapture studies (Royle et al. 2013; Sutherland et al. 2015; Fuller et al. 2016) as a model to describe movements of individuals about their home range, and also as a model for dispersal of individuals (Graves et al. 2014).

## 2 Data structure and model

Consider an idealized acoustic sampling array shown in Figure 1, which suppose is a 600 m × 600 m block of forest for which lidar measurements are available and aggregated to 20 *m*^2^ resolution showing a standardized form of average vertical vegetation density at each point (color-coded in Fig. 1). Within this landscape an array of 9 bioacoustic sensors is situated in a regular grid, and among the sensor array are located 16 experimental sources producing vocalizations that may or may not be detected at each sensor. In practice one might imagine more attenuation between a source and sensor when dense habitat (green) predominates and less attenuation in open habitats such as gaps in the forest canopy (white).

**Figure 1:**
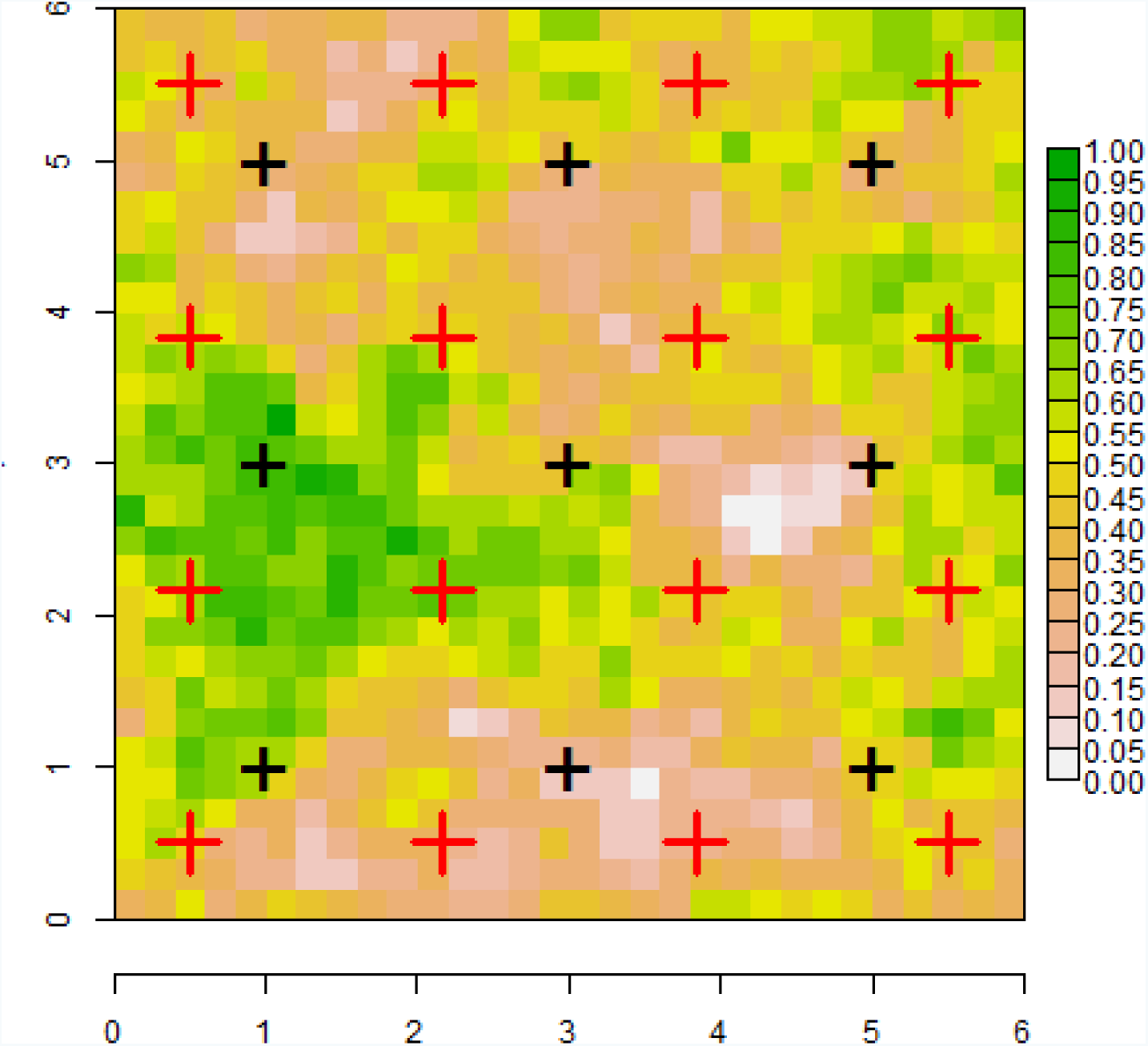
Idealized system showing an array of 9 sensors and an array of 16 experimental sources (e.g., speakers producing vocalizations of a species). Habitat structure is illustrated here as a standardize variable of vegetation density with high vertical density (green) representing dense vegetation and low vertical density (white) representing open areas.

The data from a field experiment of this sort are power or signal strength measurements, *S*_*ij*_, at each sensor having location x_*j*_, from each source *i* = 1, 2, …, *n* (*n* = 16 in this case) having location s_*i*_. The model for these observations can be formulated in terms of other signal characteristics such as time of arrival (Stevenson et al. 2015) but here for clarity I adhere to a formulation in terms of signal strength alone, although the basic ideas are the same. Following Efford et al. (2009) assume a transformation of signal strength that declines with distance *d* from the source, and assume the transformation produces a normally distributed variable such that attenuation is well approximated by the model

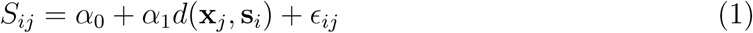

where *ε*_*ij*_ ∼ Normal(0; *σ*^2^) is noise. When the signal strength takes on positive values then the log-transformation would normally be satisfactory, and furthermore this is the natural scale when power is measured in decibels (dB). Sounds are detected when *S* exceeds a threshold *c* (Dawson and Efford 2009) which is somewhat arbitrary but should be set above the mean level of ambient noise of the system so that detections are certain to be real. The signal to noise ratio can be directly characterized from observed data (Dawson and Efford 2009). For an experimental setting where the acoustic sources are known and only detection and signal strength at each receiver are random variables, the observed data are (*y*_*ij*_, *S*_*ij*_) where *y*_*ij*_ = 1 if a signal from source *i* was detected at receiver *j* and *y*_*ij*_ = 0 if the signal was not detected, and *S*_*ij*_ > *c* is the observed signal strength (transformed as noted above). Thus, the probability of detection is *p*_*ij*_ = Pr(*S*_*ij*_ > *c*) which can be computed from the normal cumulative distribution function. When the observed signal strength is ≤ *c* it is regarded as a missing value with probability 1 − *p*_*ij*_.

Equation 1 is a basic model of sound attenuation where the attenuation of sound intensity is governed by a single parameter *α*_1_, and relates only to the Euclidean distance between source and sensor, *d*(x_*j*_, s_*i*_). Importantly, the form of this attenuation model is stationary (does not vary in space) and isotropic (it’s two-dimensional contours are circular and symmetric). Intuitively, then, this model for signal strength is probably only suitable for homogeneous environments. In what follows I propose to generalize this model by allowing for the distance *d*(x, s) to be both nonstationary and anisotropic using a non-Euclidean distance metric that, in general, depends not only on the locations of sources and sensors but also on the composition of the landscape *between* them.

### 2.1 Cost-weighted distance models

An intuitively appealing model for sound attenuation in heterogeneous environments is the cost-weighted distance (CWD) model in which attenuation is governed not by Euclidean distance but by a cost-weighted distance metric which depends on the habitat structure in the vicinity of the sensor. The cost-weighted distance can be computed for a path 
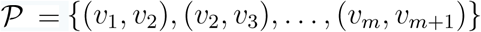
 consisting of *m* segments between any two points v_1_ and v_*m*+1_ on the landscape and it is defined by

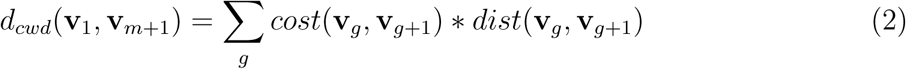

where *cost*(v_*g*_, v_*g*+1_) is a parametric function describing the cost of movement between pixels *v*_*g*_ and *v*_*g*+1_, which must be prescribed (see below) and *dist*(v_*g*_, v_*g*+1_) is the Euclidean distance between pixels. The cost-weighted distance then is the sum over all pixels along a given path connecting v_1_ and v_*m*+1_. The least-cost path (LCP) (Adriaenson et al. 2003) is the path which has minimum CWD among all possible paths connecting the points v_1_ and v_*m*+1_. In practice the cost-weighted distance between any two points and the least-cost path can be computed using the R package gdistance (van Etten 2017). Either the cost-weighted distance between points or the least-cost path can serve as an effective distance metric in models of sound attenuation, where parameter(s) of the cost function are estimated explicitly from data (see below).

The relevance of this distance metric to inference about sound attenuation arises when the cost function is parameterized in terms of the landscape structure. For example, if a covariate *z*(v) exists then one sensible function describing the cost of passing from pixel v_*g*_ to pixel v_*g*+1_ is

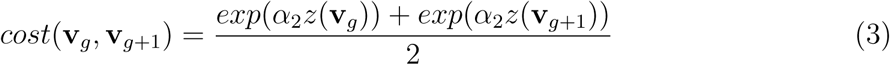

The parameter *α*_2_ represents the resistance of the covariate *z*(v) (higher values incur higher cost of transmission and *vice versa*), and it should be estimated from *observed* data on signal strength or time of arrival. I provide an estimation framework based on maximum likelihood below. To acknowledge this new distance metric in the model for sound attenuation, and that it depends on an unknown parameter *α*_2_, express the model as

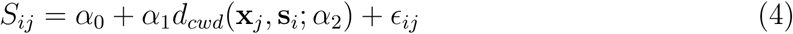

In general attenuation is frequency dependent (Wiley and Richards 1982) and thus param eters *α*_1_ and *α*_2_ should depend on species.

Obviously any number of covariates can be included in the cost function Eq. 3. Note that if *α*_2_ = 0 then the cost of transmission between any two pixels on the landscape is 1.0, and the cost-weighted distance reduces to Euclidean distance. As a practical matter we should scale any covariate *z*(v) to be in [0, 1] so that *α*_2_ can be any real number. Negative numbers imply that increasing values of the covariate facilitate sound transmission and positive values imply that increasing values of the covariate impede sound transmission. The cost-weighted distance is conveniently computed in the R package gdistance using the accCost function, and the least-cost path between any two points can be computed using the function costDistance.

To see the effect of cost weighted distance on “effective distance” Fig. 2 shows contours of effective distance (in this case the least-cost path) for different values of the resistance parameter from Eq. 3. These effective distance contours become closer together in areas of density vegetation (green) as the resistance parameter *α*_2_ increases. The basis of this as a model for sound attenuation is clear: individuals vocalizing from a location with high densities of vegetation (or other structure) between that location and the sensor should produce reduced signal strength and lowered detection probability due to sound deection, absorption and other mechanisms. In what follows I describe the model formally and demonstrate that the actual parameters governing attenuation of the sound can be explicitly estimated from experimental data on such an array.

**Figure 2:**
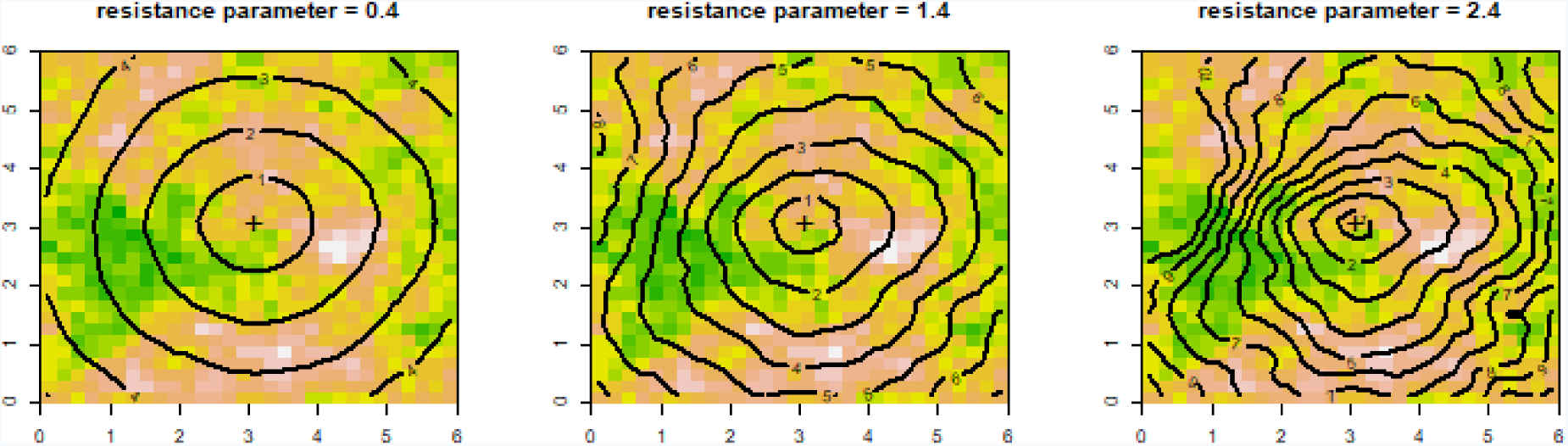
Effective distance to a sensor (shown by +) placed at (3,3) under the least-cost path model with parameter *α*_2_ = 0.4 (left), *α*_2_ = 1.4 (center) and *α*_2_ = 2.4 (right). As resistance increases, effective distance contours get closer together in response to dense structure (green).

In practice, this model could be applied in situations where relatively fine scale habitat structure data are available. For example, in a study of birds on a landscape it might be possible to obtain such data from auxiliary surveys of vegetation structure but most likely fine-scale remotely sensed data from aerial imagery, lidar or similar platforms would be ideal for this purpose. In aquatic environments attenuation is most affected by depth and sub-surface structure and in most studies of aquatic systems detailed data exist for these attributes (and others).

## 3 Likelihood Analysis

The cost-weighted distance metric described above is amenable to direct likelihood analysis from data on observed signal strength at fixed locations and with fixed sources (e.g., as in Fig. 1). The observed data from an experiment are the detection/signal strength pairs (*y*_*ij*_, *S*_*ij*_) for each source and each sensor. Recall that signal strength is truncated at some value c chosen to reflect a reasonable threshold below which signals cannot be distinguished from ambient noise. Conditional on the *J* known source locations x_*j*_, the likelihood for the data from source location s_*i*_ is

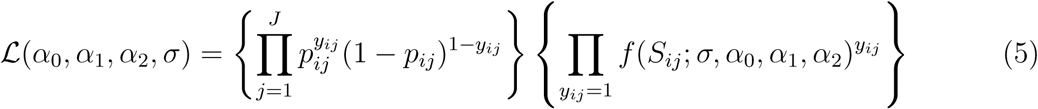

where *f*(*S*_*ij*_; *σ*, *α*_0_, *α*_1_, *α*_2_) is the normal probability density with mean *α*_0_ + *α*_1_*d*_cwd_(x_*j*_, s_*i*_; *α*_2_) and variance *σ*^2^ and the probability of detection *p*_*ij*_ = Pr(*S*_*ij*_ > *c*) which depends on the parameters of the normal distribution model for *S*_*ij*_ in Eq. 4. The likelihood can be optimized numerically using standard methods such as implemented in the R functions `nlm` or `optim` (see Appendix A).

### 3.1 Unknown source locations

The parameters of the attenuation model can be estimated from data obtained when the
sources are unknown. Of course this would be the case in any real field application of bioacoustics where animal sounds are measured. Indeed, this is precisely the situation addressed in spatial capture-recapture applications such as considered by Efford et al. (2009) and others. In this case, we have to regard the source location as a latent variable and remove it from the conditional-on-s likelihood (Eq. 5) by integrating over the planar state space (or 3-dimensional state-space in the context of aquatic systems) in the vicinity of the sensor array. One twist to the situation where the sources are unknown is the potential exists that some of the sources were not detected at all. Therefore, the likelihood has to be constructed either conditional on the event that an individual source was detected at least once (Borchers and Efford 2008) or else the possibility of *n*_0_ unobserved all-zero encounter histories must be accounted for, where *n*_0_ is then an additional parameter to be estimated (equivalently *N* = *n*_0_ + *n*_*obs*_). Indeed, *n*_0_ is the key parameter of interest in spatial capture-recapture applications. See Efford et al. (2009) for details on the likelihood construction. I provide an implementation in R of the likelihood in terms of *n*_0_ in Appendix A.

### 3.2 Computing the posterior distribution of a source

Bayes’ rule can be used to calculate the posterior distribution of an unobserved source given the pattern of detections, y, on the sensor array and the signal strengths, S. Note that the likelihood given in Eq. 5 is the joint distribution of the detection/non-detection data y_*i*_ and the signal strengths S_*i*_ *conditional* on the source location s_*i*_, say Pr(y_*i*_, S_*i*_|s_*i*_). Let Pr(s) denote the prior distribution for s, then the posterior distribution of s_*i*_ is

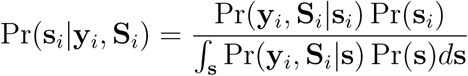

These probability distributions depend on the model parameters as in the likelihood given above but I omit that dependence to be concise. A standard assumption in spatial capture-recapture is to assume no *a priori* information about the location of a source so that Pr(s) = constant (Efford et al. 2009) in which case the posterior distribution is just standardized by the integral of the likelihood over the region in the vicinity of the sensor array. More generally, source density gradients can be accommodated by modeling explicit covariate effects in Pr(s). For example, suppose the sound sources are birds and they are likely to be using habitat preferentially, even the same habitat which is affecting sound attenuation, then we might assume

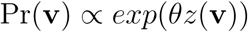

where *z*(v) is the measured habitat structure for any location v and *θ* is a parameter to be estimated.

R code for computing the posterior distribution of detected sources is given in Appendix A and I show an example in the following section.

### 3.3 Data acquisition

The model as specified here assumes that unique vocalizations can be identified and reconciled among the detectors. For example this is easily true in an experimental setting when a sound is played, in which case the sensors at which it is detected can be noted directly. Over a period of time, each individual source can be played sequentially or even replicated multiple times. In field settings when the source location is unknown then a specific source encounter history has to be reconciled in a sense manually. But in practice can be done unambiguously in many practical settings if the density of sources is not too high (Dawson and Efford 2009). In the field (sampling real birds), an individual might make many calls during a particular time interval and these are treated as distinct sources.

## 4 Illustration

Using the experimental sensor array shown in Fig. 1 I simulated some data under the model for log-signal strength with *α*_0_ = 0, *α*_1_ = 1.0 and *σ* = 0.50. Therefore,

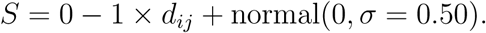

I used a threshold of detection of *c* = −3. Moreover, the least-cost path distance model of attenuation was used with *α*_2_ = 2.0 to model attenuation through the heterogeneous habitat shown in Fig. 1. These parameter settings produce an average of 20.8 total captures of 13.8 individuals on the array of 9 sensors. A particular realization is shown in Fig. 3 which shows the pattern of detections of the 16 sources. In particular, lines are connected between each source and the sensor(s) at which it was detected. Three of the sources were not detected at all, 6 were detected once, 6 were detected twice, and one source three times. The MLEs for the model parameters for this single realization are 
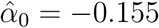
, 
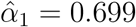
, 
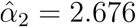
 and 
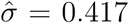
. In general, a very high level of precision is possible for this relatively low level of detections in an experimental setting when the source locations are known. For example, 100 realizations of this situation produce an MLE of *α*_2_ having mean 1.96 (recall truth = 2.0) and standard error 0.398. The R script for simulating data and fitting the model is given in Appendix A.

**Figure 3:**
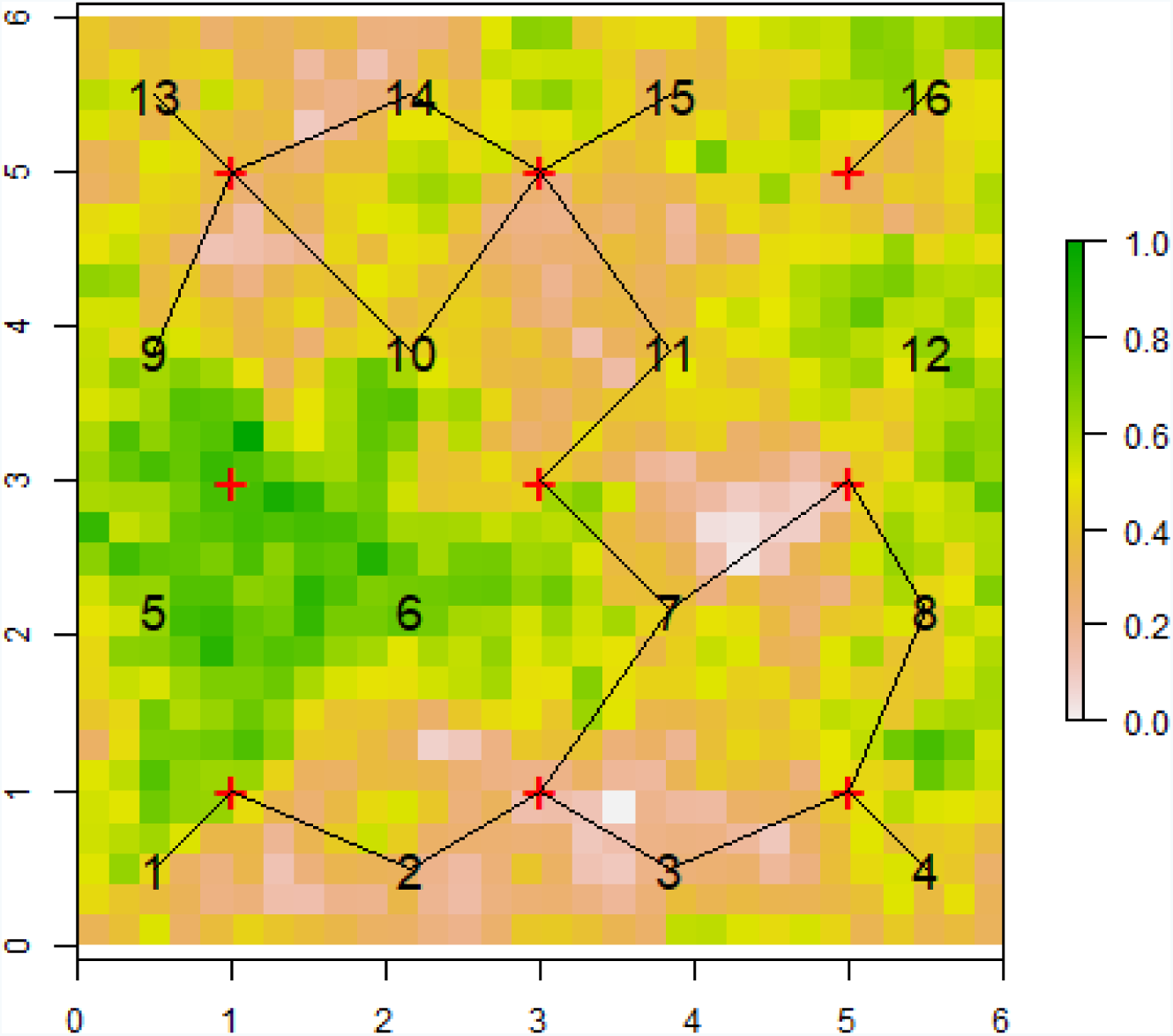
A single realization of a bioacoustic experiment to measure non-Euclidean attenuation in the form of the least-cost path model. For this simulated realization, detection frequencies of the 16 sources as follows: (1, 2, 2, 1, 0, 0, 3, 2, 1, 2, 2, 0, 1, 2, 1, 1). Each source is connected to the sensors at which it was detected.

Maximum likelihood estimation for this experimental system is much less effective when the source locations s are *unknown*, in which case the MLEs can be considerably biased (see Appendix A). Instead, a much larger array is required, or a denser population of sources is needed in order to generate sufficient encounters. This is consistent with what is known in spatial capture-recapture studies; see for example Efford and Fewster (2013), Sun et al. (2014) and chapter 10 in Royle et al. (2014)). Nevertheless, it is possible to localize the unknown sources using the general likelihood formulation based on the marginal likelihood. For the same simulated data set shown in Fig. 3 I produced the estimated posterior distribution of the unknown source location of 4 sources (Fig. 4) captured between 1 and 3 times each. We see that the estimated posterior distributions are in the vicinity of the true source locations, modified by the observed encounter history (the data set is generated using the random number seed noted in Appendix A).

**Figure 4:**
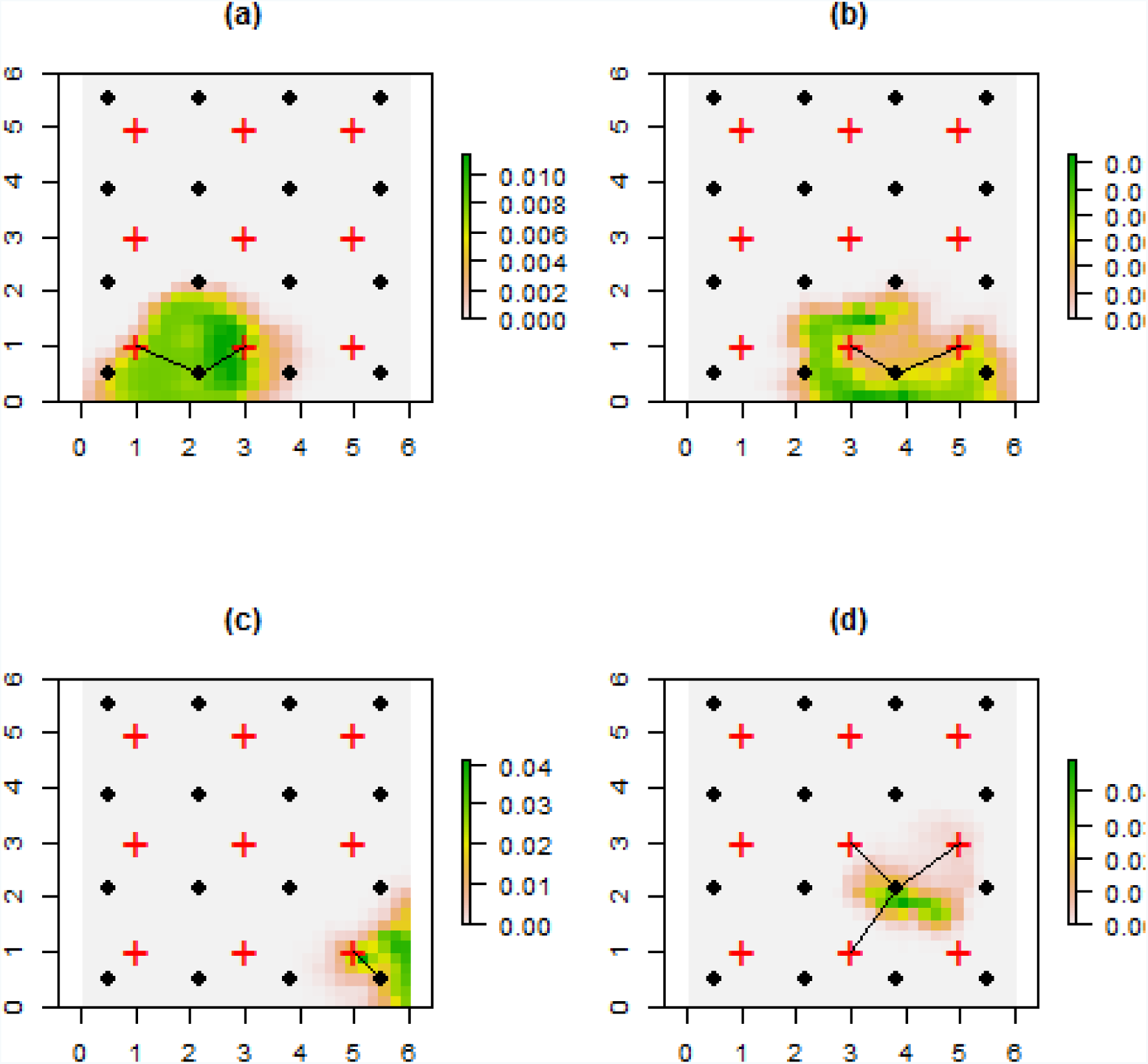
Estimated localizations of 4 unknown sources characterized by the estimated
posterior distribution of the source location obtained by plugging-in the MLEs obtained by maximizing the likelihood for the observed detection/non-detection and signal strength data. Because these are posterior distributions the sum over all pixels equals 1.0. For each of the 4 sources shown here the true location of the source is connected to the sensors at which it was detected by lines.

## 5 Discussion

With the rapid and expanding adoption of acoustic monitoring technology, the ability to understand sound attenuation along environmental gradients will become increasingly important (Kessel et al. 2013). In this paper I suggested a flexible framework for modeling sound attenuation in heterogeneous environments. This framework has two direct applications. First, it can lead to improved inferences about source locations (“localization”) which is important in many applications, especially acoustic telemetry. Second, it allows investigators to better understand how bioacoustic methods work under field conditions in heterogeneous environments by enabling *in situ* inference about factors that inuence attenuation from field data. The method is sufficiently flexible that it can be used with acoustic telemetry data, as well as encounter history data used in acoustic SCR applications (Efford et al. 2009; Stevenson et al. 2015; Kidney et al. 2016). I formulated the model here in terms of “signal strength” data (e.g., sound intensity measured in decibels) but the idea applies directly when time of arrival data are available. For such data, the localization model also involves a distance function (Stevenson et al. 2015) which might be replaced by cost-weighted or least-cost path distance with parameters to be estimated. In addition, the basic ideas apply directly to classical distance sampling methods (Buckland et al. 2001), where the Euclidean distance metric in the distance sampling likelihood can be replaced by cost-weighted distance. In the case of distance sampling, in most applications, the model would provide a description of visual obstruction and not wave attenuation.

The ability to develop explicit models of sound attenuation has important sampling design implications. In heterogeneous environments, the detection range of sensors depends on environmental characteristics (Kessel et al. 2013; Selby et al. 2016) and thus this critical parameter is both nonstationary and anisotropic. Therefore, the optimal spacing of sensors in an array must be variable in response to the underlying environmental heterogeneity. The problem of array design is analogous to the design of camera trap studies (e.g., Royle et al. 2014; ch. 10), where arrays can be constructed so as to maximize the probability of detection, optimize criteria based on the variance of estimators of parameters of interest, or maximize the precision of the localization. Using the non-Euclidean model of effective distance suggested in this paper, one could obtain estimates of the parameter *α*_2_ from an experimental or observational study and then use that estimate to improve the design of bioacoustic monitoring arrays. In practice, having multiple sensors in a given experimental array, with known source locations, such as shown in Fig. 1, is not necessary. One could obtain suitable estimates of model parameters with a single sensor and replicated source emissions. However, in field applications of localization or density estimation (such as Dawson and Efford 2009) multiple sensors are required. In general, estimation of model parameters is challenging when source locations are unknown and effective estimation might require a a large array of sensors and a large sample size of detections. As such, in practice, one might first consider experimental analysis of sound attenuation models with known sources in order to obtain precise estimates of parameters of the effective distance model which may then be treated as fixed in subsequent analyses focused on localization or density estimation.

An obvious extension of the framework proposed here is to consider alternative non-Euclidean distance metrics. For example, one obvious alternative is to substitute resistance distance (McRae 1996) in place of cost weighted distance. Resistance distance is based on an analogy between discrete landscapes (characterized by a raster of pixels) and electrical circuits. Resistance distance between two nodes is the effective resistance between them, which depends on the resistance between each node and the number of pathways in the circuit. I think both cost weighted distance (or least-cost path) and resistance distance offer useful descriptions of sound attenuation in heterogeneous landscapes.

One limitation of the proposed approach is that the cost-weighted distance model underlying least-cost path is a phenomenological model. It describes the phenomenon of attenuation in response to measurable covariates but does not explicitly embody elements of sound dynamics such as reverberation, absorption and reflection. Rather, it models their total effect as measured by the apparent relative distance between points as measured by observed signal strength and pattern of detections. This may not be a severe limitation in biological applications of acoustic monitoring where interest is usually in the end use of the data for detection, localization or similar objectives and not directly in the processes contributing to sound dynamics of a particular system.

Application of any non-Euclidean effective distance model depends on the availability of environmental or habitat information at a suitable scale to be relevant to sound dynamics. While small scale habitat data are not always collected in field studies using acoustic sampling or distance sampling, it seems likely that such data will be collected more frequently in the future with the increasing availability of lidar technology (Zolkos et al. 2013, He et al. 2015) and other remote sensing platforms such as drones (Martin et al. 2012, Christie et al. 2016).

## Acknowledgements

The author thanks Cathleen Balantic, Danielle Rappaport, Andrew MacLaren and Suresh Sethi for helpful comments and suggestions on this manuscript. Any use of trade, product, or firm names is for descriptive purposes only and does not imply endorsement by the U.S. Government.

## Appendix A

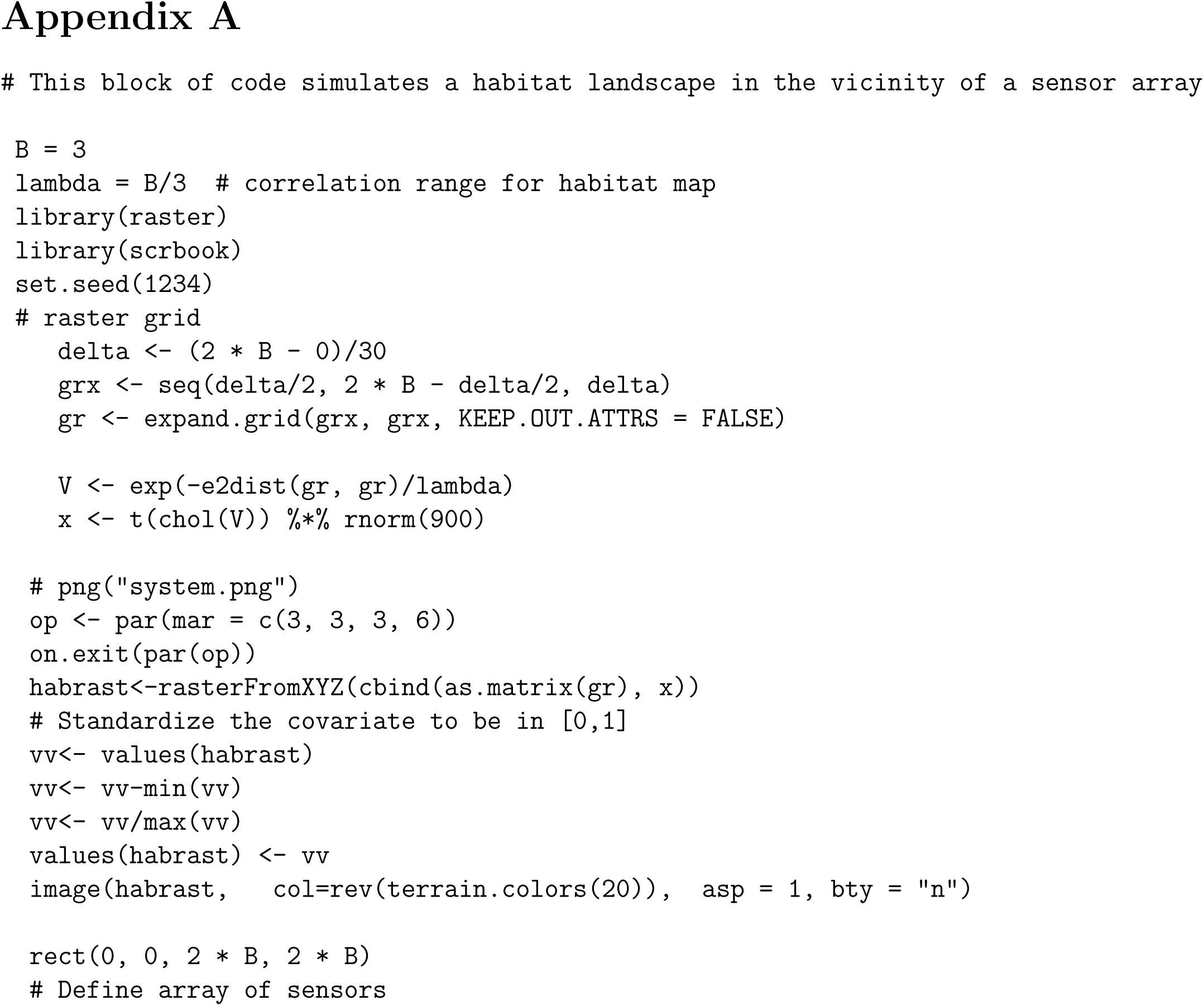
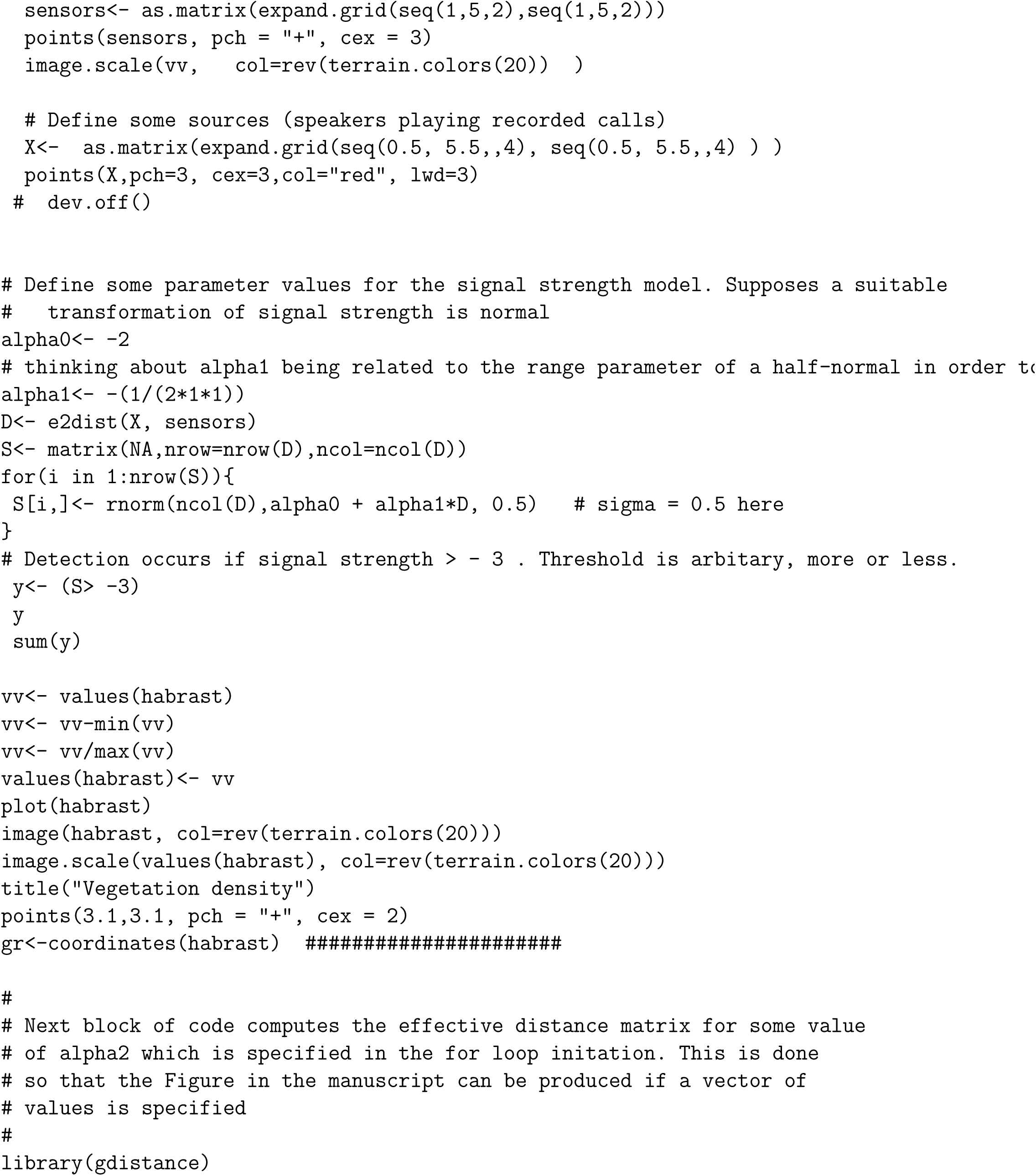
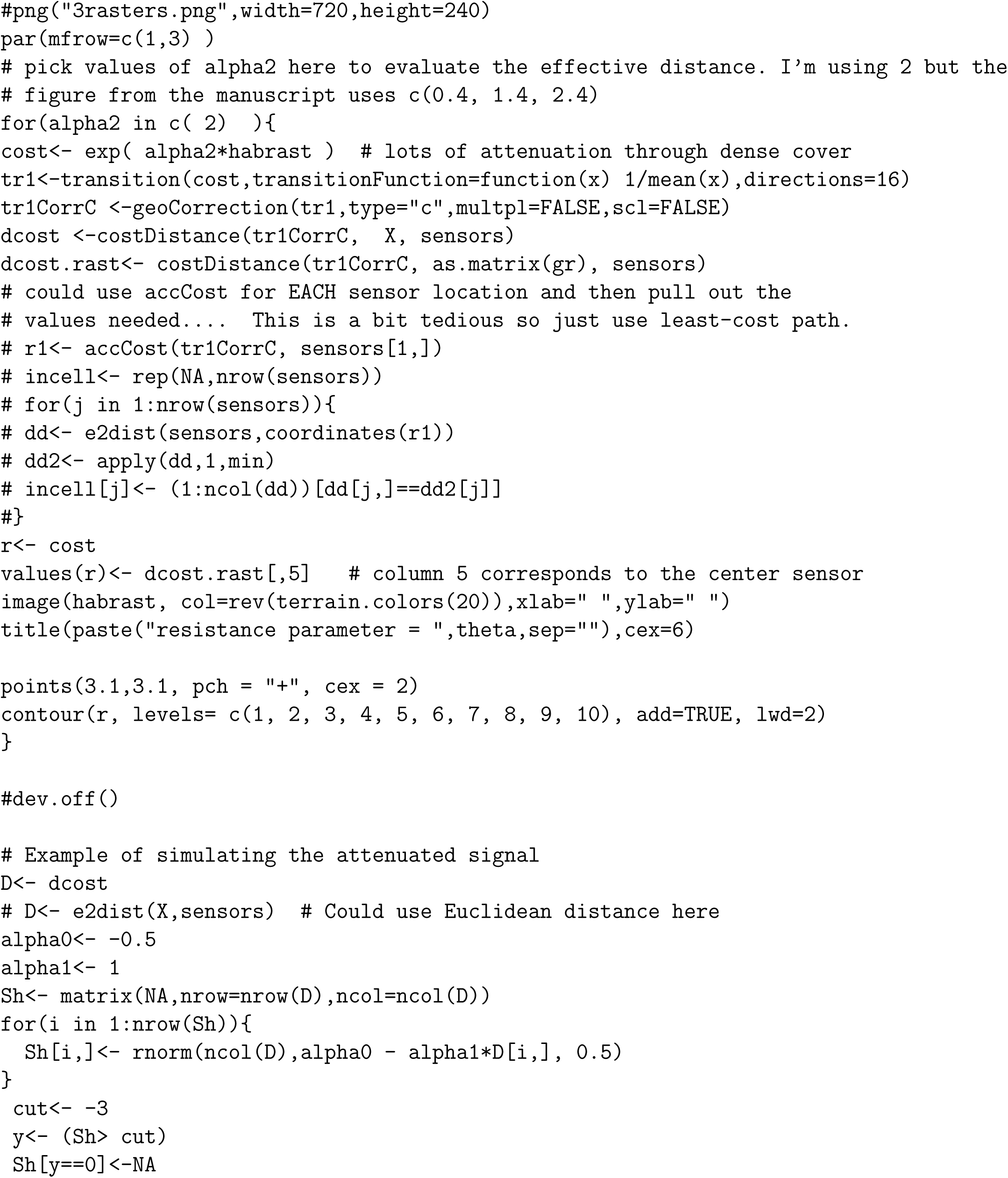
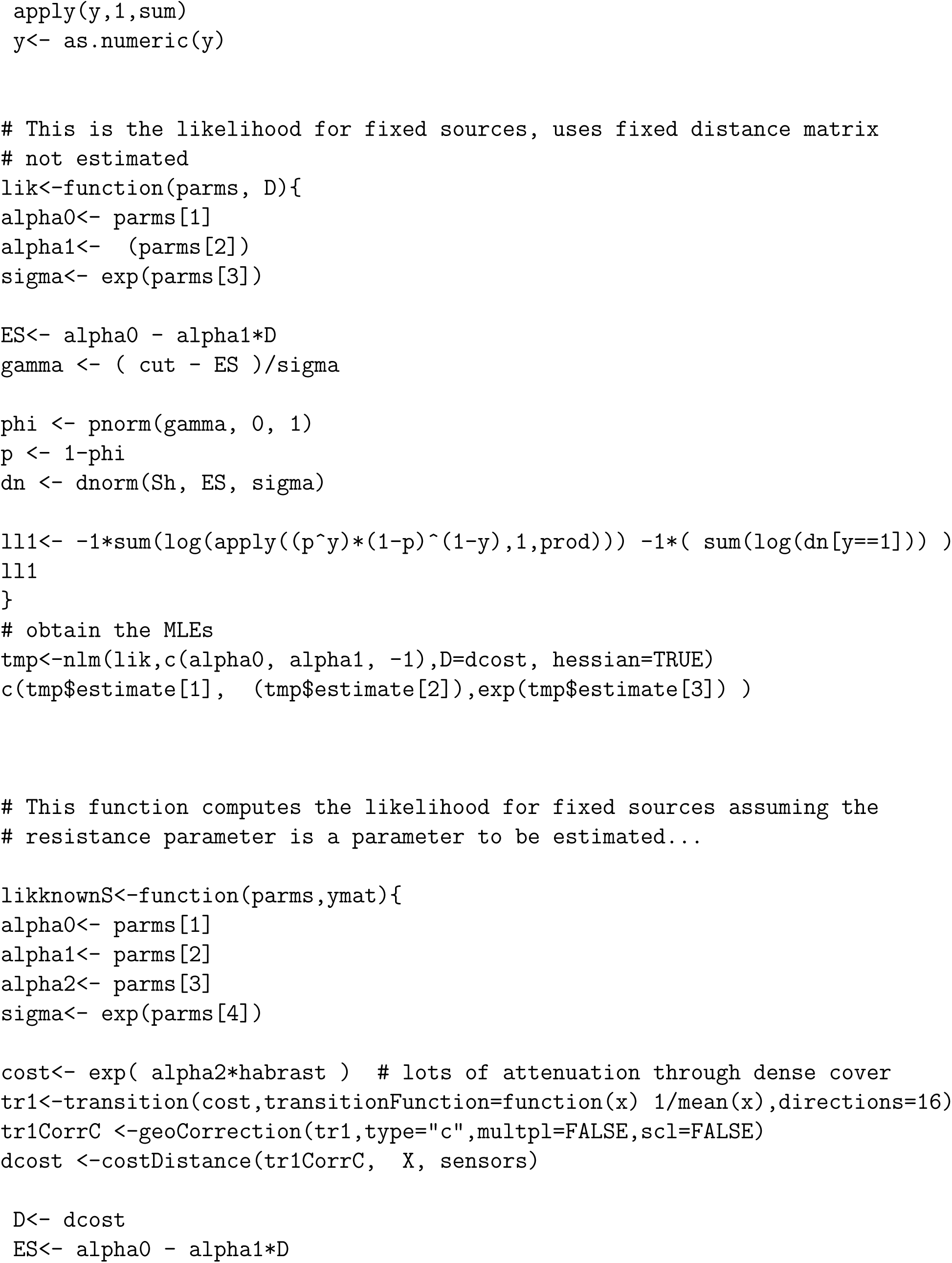
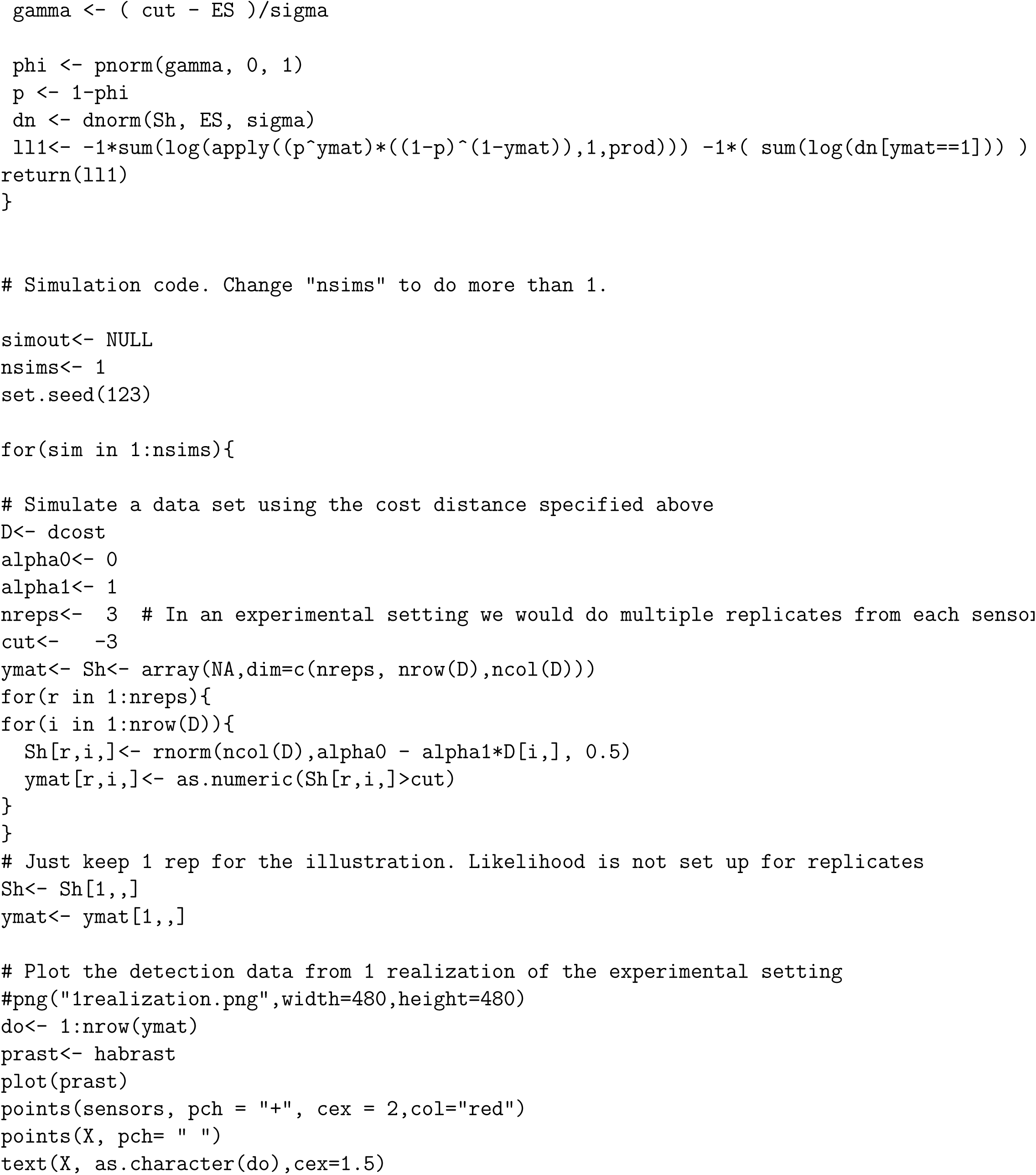
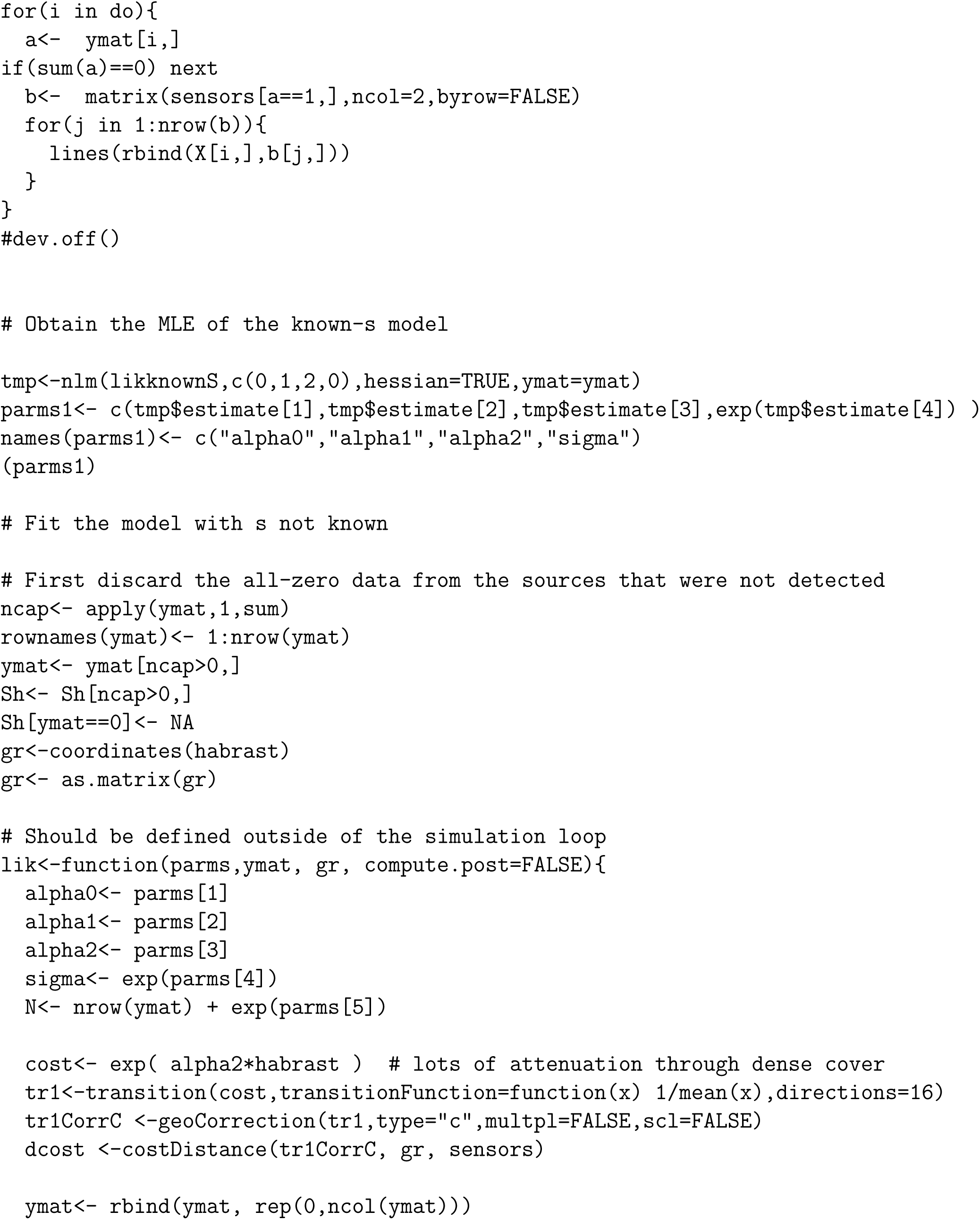
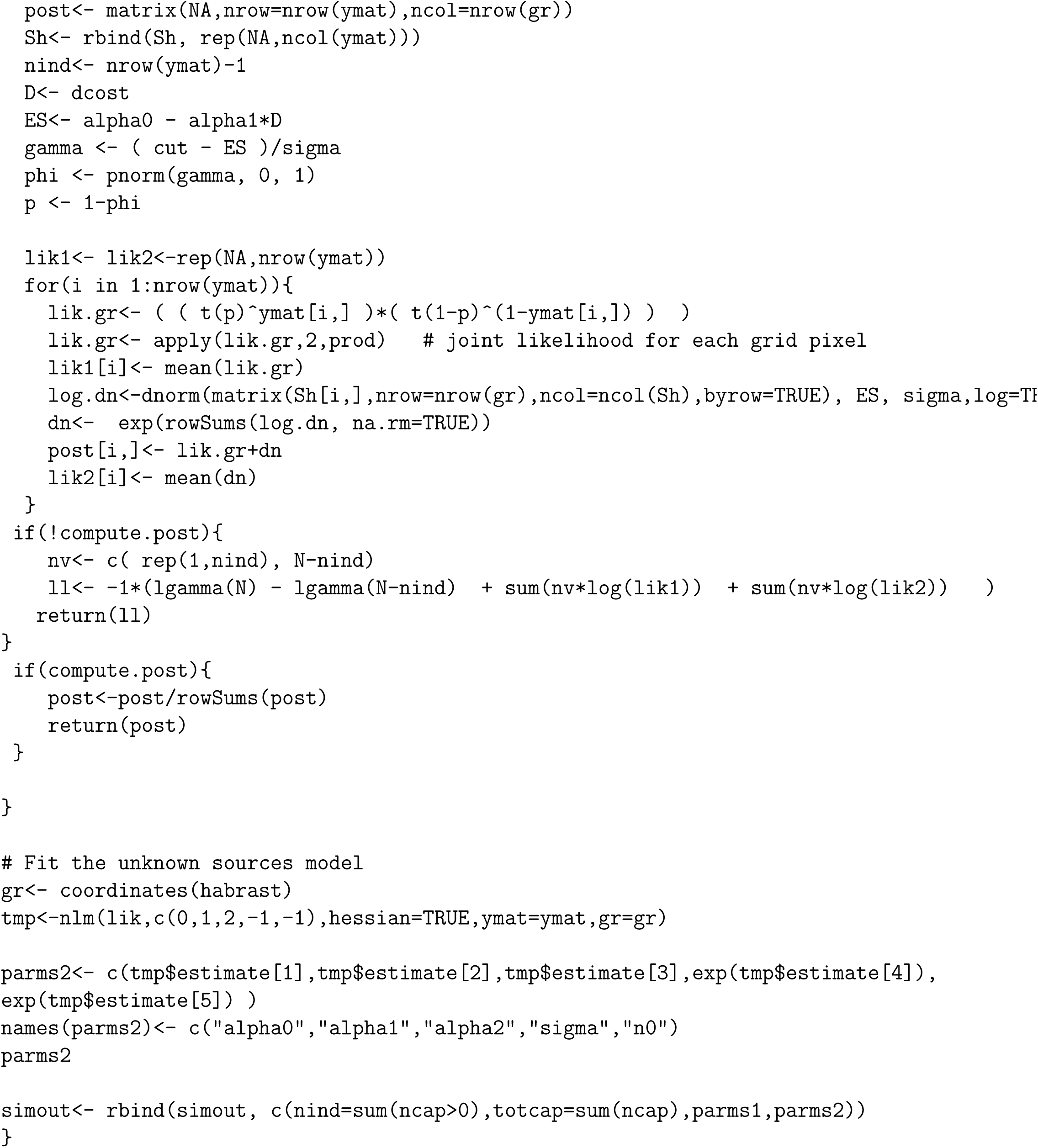
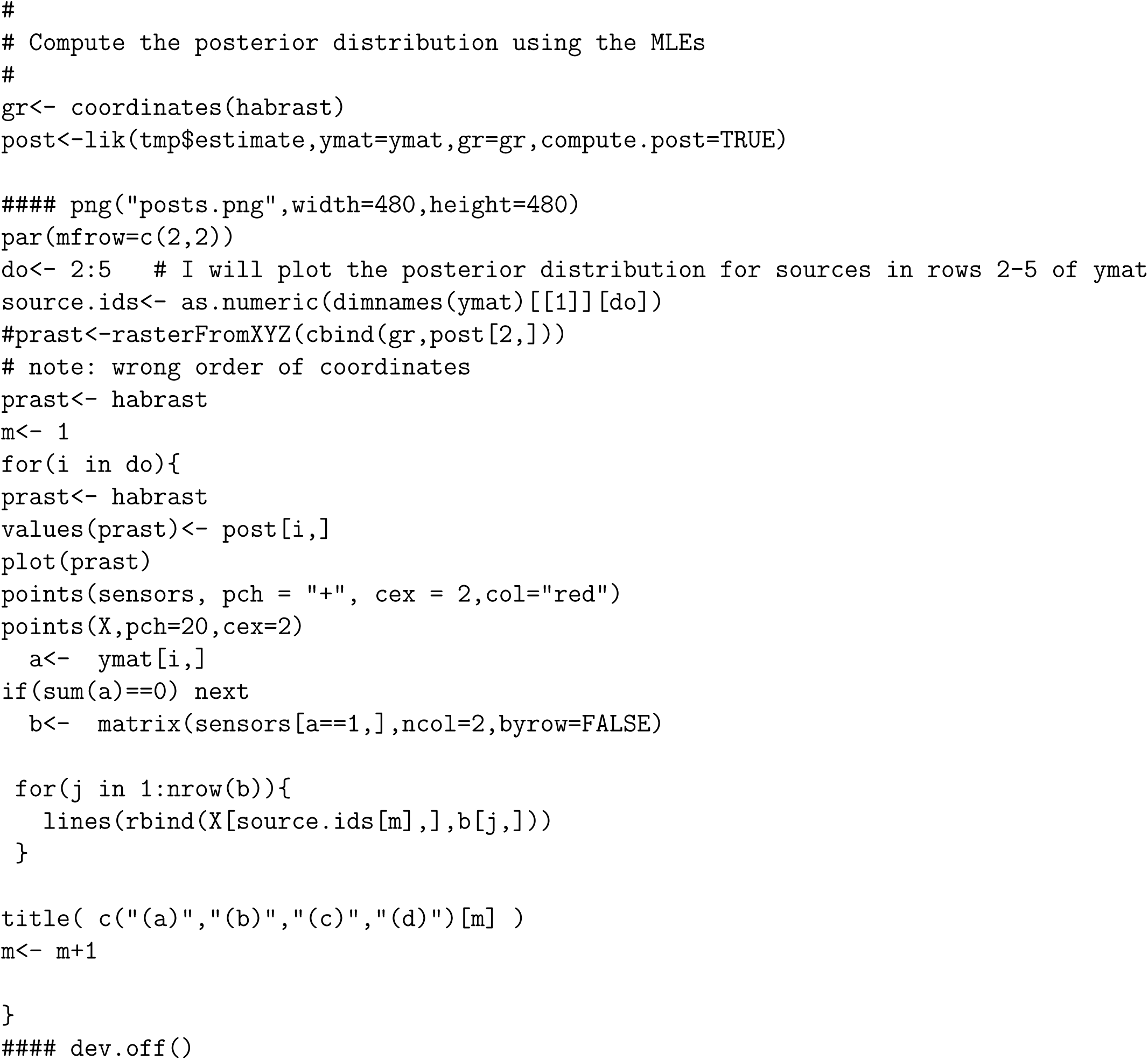

## Footnotes

1 e.g., see https://en.wikipedia.org/wiki/Acoustic_attenuation accessed 12/20/2016.

